# Comparative Genomics and Environmental Distribution of Large dsDNA viruses in the family *Asfarviridae*

**DOI:** 10.1101/2021.01.29.428683

**Authors:** Sangita Karki, Mohammad Moniruzzaman, Frank O. Aylward

**Affiliations:** Department of Biological Sciences, Virginia Tech, Blacksburg, VA, 24061, USA

**Keywords:** *Asfarviridae*, NCLDV, Megaviridae, eukaryotic viruses, *Nucleocytoviricota*

## Abstract

The *Asfarviridae* is a family of Nucleo-Cytoplasmic Large DNA Viruses (NCLDV) of which African swine fever virus (ASFV) is the most well-characterized. Recently the discovery of several *Asfarviridae* members other than ASFV has suggested that this family represents a diverse and cosmopolitan group of viruses, but the genomics and distribution of this family have not been studied in detail. To this end we analyzed five complete genomes and 35 metagenome-assembled genomes (MAGs) of viruses from this family to shed light on their evolutionary relationships and environmental distribution. The Asfarvirus MAGs derive from diverse marine, freshwater, and terrestrial habitats, underscoring the broad environmental distribution of this family. We present phylogenetic analyses using conserved marker genes and whole-genome comparison of pairwise average amino acid identity values, revealing a high level of genomic divergence across disparate Asfarviruses. Further, we found that *Asfarviridae* genomes encode genes with diverse predicted metabolic roles and detectable sequence homology to proteins in bacteria, archaea, and different eukaryotes, highlighting the genomic chimerism that is a salient feature of NCLDV. Our read mapping from Tara oceans metagenomic data also revealed that three *Asfarviridae* MAGs were present in multiple marine samples, indicating that they are widespread in the ocean. In one of these MAGs we identified four marker genes with >95% amino acid identity to genes sequenced from a virus that infects the dinoflagellate *Heterocapsa circularisquama* (HcDNAV). This suggests a potential host for this MAG, which would thereby represent a near-complete genome of a dinoflagellate-infecting giant virus. Together, these results show that *Asfarviridae* are ubiquitous, comprise similar sequence divergence as other NCLDV families, and include several members that are widespread in the ocean and potentially infect ecologically important protists.

## 1 Introduction

The Nucleo-Cytoplasmic Large DNA Viruses (NCLDVs), also called *Nucleocytoviricota,* comprise a phylum of dsDNA viruses that infect diverse eukaryotes (Van Etten et al., 2010b; Koonin et al., 2020). NCLDVs include the largest viruses known, both in terms of virion size and genome length, and genomes within this group often contain genes involved in metabolic pathways that are otherwise present only in cellular lineages (Fischer et al., 2010; Van Etten et al., 2010b; Schvarcz and Steward, 2018; Moniruzzaman et al., 2020a). Some families of NCLDV such as the *Poxviridae, Asfarviridae, Iridoviridae*, and *Phycodnaviridae* have been studied for decades, while others, such as the *Pandoraviridae, Mimiviridae,* and *Marseilleviridae,* have been discovered relatively recently (Raoult et al., 2004; Boyer et al., 2009; Philippe et al., 2013; Abergel et al., 2015). Although amoebae have been used as an effective system to cultivate many recently-discovered NCLDV, recent cultivation-independent studies have discovered a wide range of these viruses in diverse environments, suggesting that uncultivated members of this viral phylum are ubiquitous in the biosphere (Monier et al., 2008; Hingamp et al., 2013; Bäckström et al., 2019; Endo et al., 2020; Moniruzzaman et al., 2020a; Schulz et al., 2020). Given the notable complexity of NCLDVs and their cosmopolitan distribution, there is a need to better understand their genomic diversity and biogeography.

The *Asfarviridae* is a family of NCLDVs for which the most well-studied member is the African Swine Fever Virus (ASFV), an emerging pathogen that was first discovered in 1921 (Montgomery and Eustace Montgomery, 1921). Although ASFV has been extensively studied due to its high mortality and subsequent economic toll on livestock production, other viruses within the same family have remained relatively underexplored, and until recently the vertebrate pathogen ASFV was the only known member of the *Asfarviridae* family. In 2009 a virus infecting the marine dinoflagellate *Heterocapsa circularisquama* was cultivated, and partial sequencing of the DNA polymerase type B and MutS genes revealed that the virus likely belonged to the *Asfarviridae* (Ogata et al., 2009). Furthermore, a new amoeba virus, Faustovirus and other isolates of amoeba-infecting Asfarviruses that clustered with the *Asfarviridae* have also been reported (Reteno et al., 2015; Benamar et al., 2016). Using amoeba as a cell support, two other circular dsDNA virus isolates, Kaumoebavirus and Pacmanvirus that clustered with *Asfarviridae* were obtained (Bajrai et al., 2016; Andreani et al., 2017). Lastly, a culture independent study in early 2020 reported Asfar-like virus (AbalV) causing mass mortality in abalone (Matsuyama et al., 2020). Together, these studies have begun to show that the *Asfarviridae* are likely a diverse family of NCLDV that are globally distributed and infect both protist and metazoan hosts.

Recently, two studies (Moniruzzaman et al., 2020a; Schulz et al., 2020) reported numerous new metagenome-assembled genomes (MAGs) of NCLDV, some of which have phylogenetic affinity with the *Asfarviridae* family. However, the genomic characteristics of these MAGs have not been studied in detail. In this study, we leveraged five previously available Asfarvirus genomes and 35 new Asfarvirus MAGs to perform comparative genomic and biogeographic analysis of the *Asfarviridae* family and provide an assessment of the scale of Asfarvirus diversity in the environment. We assess the phylogenetic relationship of these new MAGs and previously discovered Asfarviruses to explore their evolutionary relationships, and we identify the potential evolutionary origins of the *Asfarviridae* genomic repertoires. We also report numerous genes encoding for different metabolic activities including central amino acid metabolism, nutrient homeostasis, and host infection. Moreover, we assess the distribution of marine Asfarvirus genomes in the ocean, and we identified high sequence similarity between one marine Asfarvirus MAG to only few proteins sequences available from a virus known to infect the dinoflagellate *Heterocapsa circularisquama,* suggesting a potential host for this MAG. Our findings reveal that the *Asfarviridae* members are widespread in the ocean and potentially have roles in biogeochemical cycling through infection of key protist lineages.

## 2 Materials and Methods

### 2.1 Comparative analysis and Protein annotation

For this study, we analyzed 35 Asfarvirus MAGs generated in two previous studies (Moniruzzaman et al., 2020a; Schulz et al., 2020) and complete genomes of five Asfarviruses (Reteno et al., 2015; Silva et al., 2015; Bajrai et al., 2016; Andreani et al., 2017; Matsuyama et al., 2020). MAGs were quality-checked using ViralRecall v. 2.0 (default parameters), with results manually inspected to ensure that no large non-NCLDV contigs were present (Aylward and Moniruzzaman, 2021). We used Seqkit v0.12.0 (Shen et al., 2016) for FASTA/Q file manipulation to generate the statistics of the genomes and proteins. To predict protein and search for tRNA genes, we used Prodigal V2.6.3 (Hyatt et al., 2010) and ARAGORN v1.2.38 (Laslett, 2004) respectively with default parameters. For the sequence similarity search, we used BLASTp against the NCBI reference sequence (RefSeq) database, version 92 (O'Leary et al., 2016). An E-value threshold of 1e-3 was used, and maximum target sequence was set to 1 so that only the best alignment for each query protein is provided. Functional annotation of predicted proteins was done using hmmsearch (parameter -E 1e-5) in HMMER v3.3 (Eddy, 2011) against the EggNOG v.5 database (Huerta-Cepas et al., 2016) to assess the potential function of MAG-encoded proteins, and the best hits for each protein were recorded.

We calculated protein-level orthologous groups (OGs) shared between all genomes analyzed in this study using the Proteinortho tool version 6.0.14 (Lechner et al., 2011) with default parameters. The resulting matrix for the orthologous genes was used for the bipartite network analysis. A bipartite network for the 35 MAGs along with their reference genomes were constructed using igraph (Csardi and Nepusz, 2006), selected members of *Poxviridae* were used as an outgroup. The network consisted of two node types, one for genomes and one for orthologous groups. OGs that were present in at least one genome were analyzed. A Fruchterman-Reingold layout with 10000 iteration was used for visualization purposes.

To assess the genomic diversity between Asfarviruses, we calculated Amino acid Identity (AAI) using the python script available at https://github.com/faylward/lastp_aai. This script uses LAST to detect bi-directional best hits to find the pairwise identity of orthologous proteins (Kiełbasa et al., 2011). The results were visualized using the gplots package (Warnes et al., 2020) in the R environment.

In order to assess the sequence similarity, the raw metagenomic reads from TARA ocean samples described previously (Sunagawa et al., 2015) were downloaded from the NCBI SRA database, and forward Illumina reads were mapped against the selected genomes using LAST (Kiełbasa et al., 2011) with default parameters. The results were visualized with fragment recruitment plots using the ggplot2 package (Wickham, 2009) in the R environment.

### 2.2 Phylogenetic reconstruction

To generate the phylogenetic tree, we analyzed 35 MAGs and five reference genomes along with selected members of the *Poxviridae* as an outgroup. We used five marker genes: Major Capsid Protein (MCP), Superfamily II helicase (SFII), Virus-like transcription factor (VLTF3), DNA Polymerase B (PolB), and packaging ATPase (A32), that are previously shown to be useful and used for phylogenetic analysis of NCLDV MAGs (Yutin et al., 2009; Moniruzzaman et al., 2020a). We used a python script that ensures the proteins have hit to the same HMM and merge those proteins together to form non overlapping alignments with HMM (available at github.com/faylward/ncldv_markersearch), also previously described (Moniruzzaman et al., 2020a). We used Clustal Omega v1.2.4 (Sievers et al., 2011) for alignment, trimAl v1.4.rev15 (Capella-Gutierrez et al., 2009) for alignment trimming (parameter -gt 0.1). We used IQ-TREE v. 1.6.12 (Minh et al., 2020) with the “-m TEST” model finder option (Kalyaanamoorthy et al., 2017) that identified VT+F+I+G4 as the best- fit model and 1000 ultrafast bootstrap (Hoang et al., 2018) to reconstruct a maximum likelihood phylogenetic tree. Finally, we visualized the resulting phylogenetic tree using Interactive Tree of Life (iTOL) (Letunic and Bork, 2019).

Another phylogenetic tree was built using only PolB as a marker gene with the methods described previously. We did this because we observed that one NCLDV MAG (ERX552270.16) contained a PolB sequence with >98% amino acid identity to the PolB sequenced from the *Heterocapsa circularisquama* virus HcDNAV (Ogata et al., 2009) (as ascertained using BLASTP), and we wanted to confirm that these sequences clustered together. The complete proteome of HcDNAV is not available, and so inclusion of this virus in the multi-locus tree was therefore not possible.

## 3 Results and Discussion

### 3.1 Asfarvirus genome statistics

The Asfarvirus MAG assembly sizes ranged from 120 kbp (SRX802982.1) to 580.8 kbp (GVMAG-S-3300009702-144). Among the 35 MAGs, 17 had all five core genes used for phylogenetic analysis (A32, PolB, MCP, SFII, VLTF3) while the rest of the genomes were missing only one core gene, including three MAGs in which the highly conserved PolB marker was not identified. This suggests that the MAGs are generally high quality, although the absence of some marker genes suggests that some are only nearly complete and that MAG assembly sizes are underestimates of the complete genome sizes. The % G+C content for the new MAGs ranged from 17 % to 60%, while those of reference viruses ranged from 31 to 45%. The ARAGORN software predicted three tRNA genes (Leu, Ile, and Asn) for ERX552270.16, one Ile-tRNA gene for GVMAG-M-3300013133-40, GVMAG-M-3300023174-161, GVMAG-M-3300027793-10, GVMAG-S-3300005056-23, and GVMAG-S-3300010160-169, and one Arg- tRNA gene for SRX319065.14. One tRNA gene (Ile) was also predicted in reference virus- Pacmanvirus as described previously (Andreani et al., 2017). The complete statistics for the MAGs are provided in Table 1.

**Table 1.**
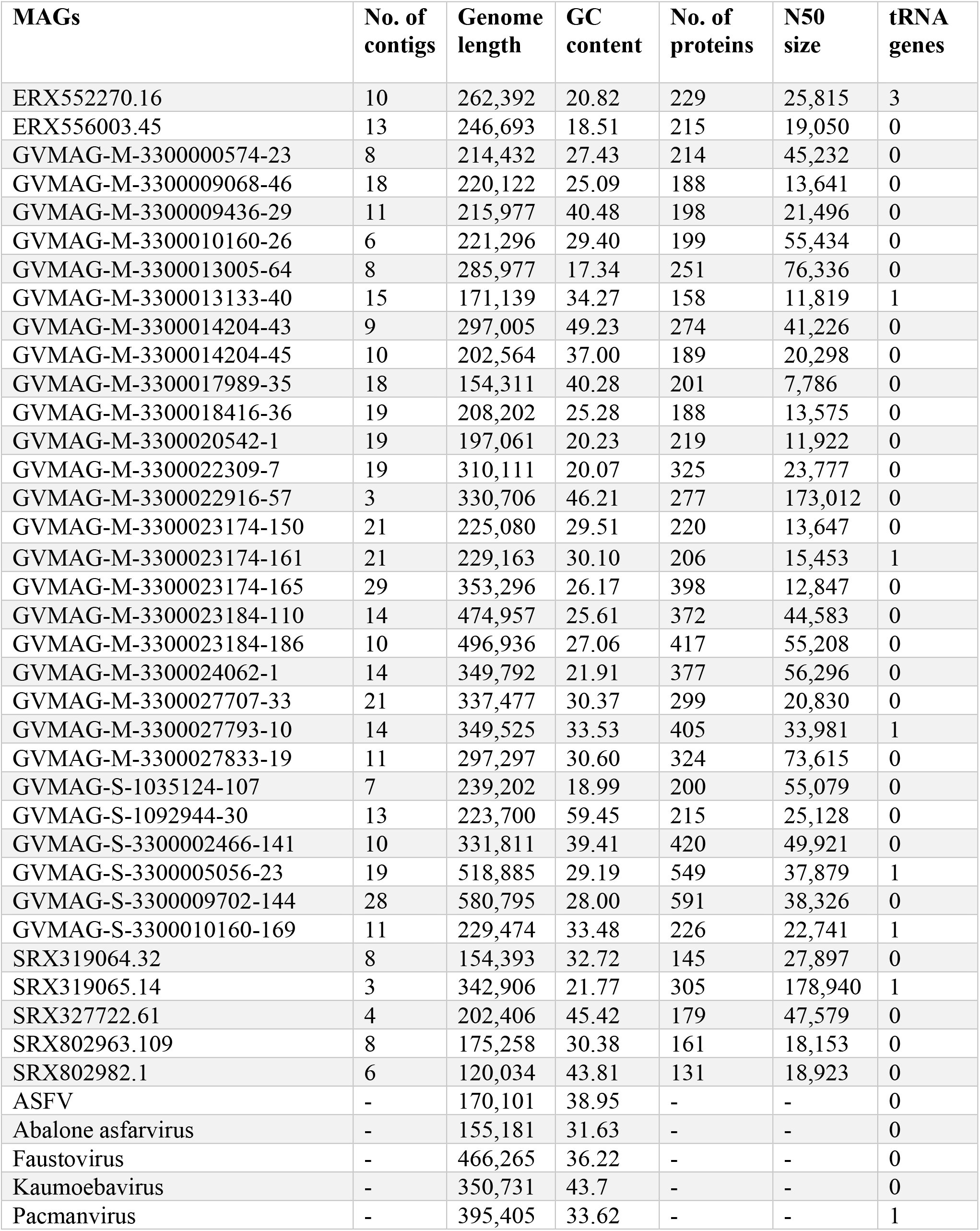
General statistics of the five Asfarvirus genomes and 35 viral MAGs

### 3.2 Phylogenetic relationship between the Asfarviruses

To assess the phylogenetic diversity and evolutionary relationships of the new MAGs, we constructed a phylogenetic tree based on alignment of the five conserved marker genes. These marker genes have been previously described to be highly conserved in the NCLDVs (Yutin et al., 2009; Moniruzzaman et al., 2020a). The phylogenetic analysis revealed that although the Asfarvirus MAGs formed clades with the five reference genomes (ASFV, Abalone asfarvirus, Kaumoebavirus, Faustovirus, and Pacmanvirus) in some cases, overall, the new MAGs had deep branches and were not closely related to cultivated viruses. The numerous deep-branching lineages in the tree underscores the high level of phylogenetic divergence between different Asfarviruses. The new MAGs were isolated from different environments, including freshwater (18), marine (12), landfill (2), non-marine saline lake (2), and mine tailing samples (1), highlighting their broad distribution. Clustering of the isolates according to the environment was also apparent in the phylogenetic tree, with several clades found only in marine or freshwater environments (Figure 1). This suggests that the broad habitat preference of many Asfarviruses may be conserved across some clades.

**Figure 1.**
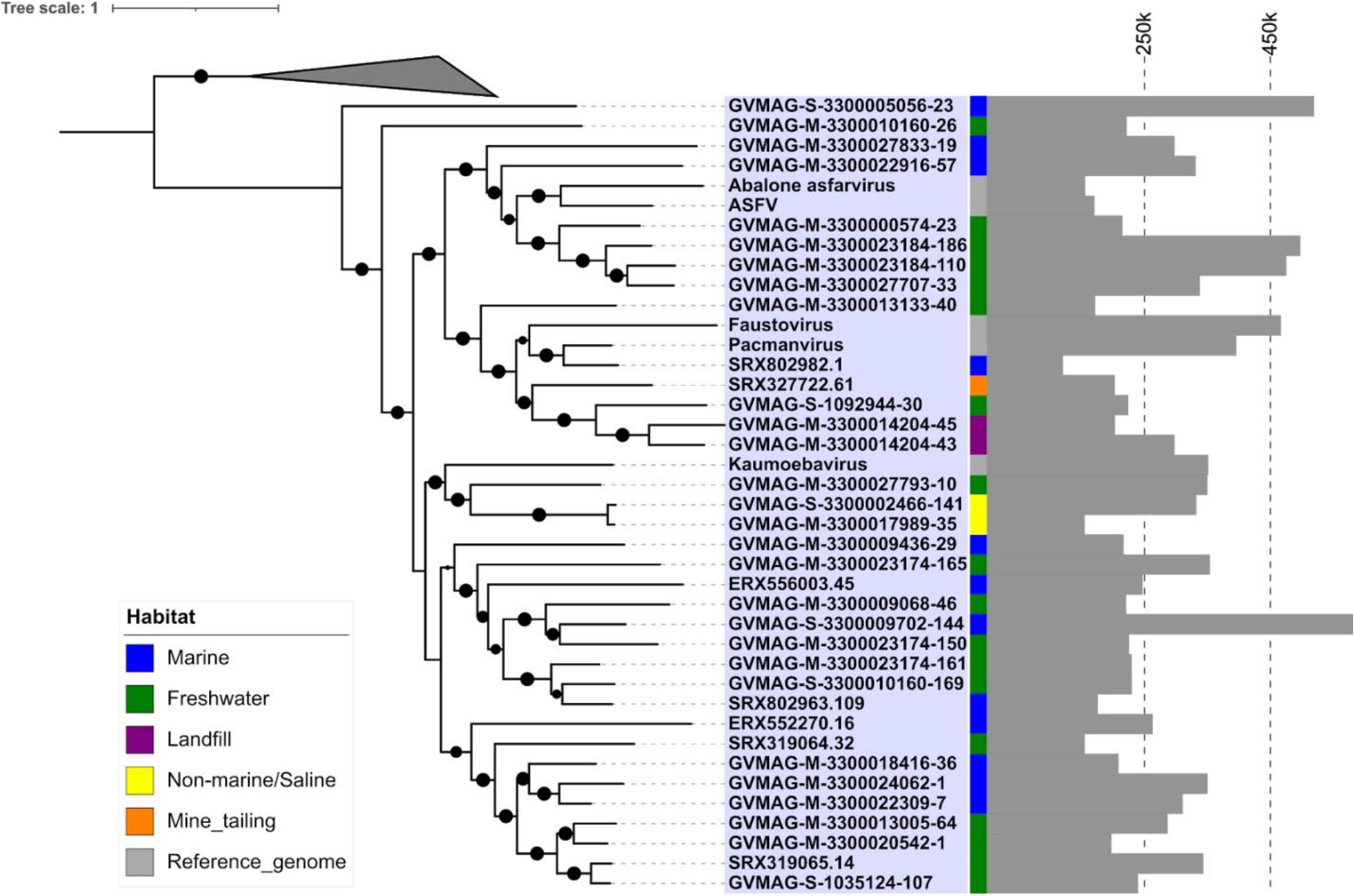
Phylogenetic tree based on five conserved marker genes. (The inner strip represents the habitat while the bar chart with scale represents the genome size of the MAGs in bp). The size of the black dot represents the bootstrap values. Only bootstrap values greater than 0.5 are shown.

The MAG GVMAG-S-3300005056-23 was the most basal-branching Asfarvirus genome. We compared the proteins encoded in this genome to the NCBI RefSeq database and found that 13 had best hits to Poxviruses (compared to at most 4 in the other Asfarvirus MAGs), while 37 proteins had best hits to Asfarvirus genomes in this database (Supplementary Data 1). Together with its basal placement in our phylogeny, these results suggest that GVMAG-S-3300005056-23 is either a basal branching Asfarvirus or possibly even a member of a new family of NCLDV. We chose to use Poxviruses to root our phylogeny because this family is often considered to be most closely related to the *Asfarviridae* (Iyer et al., 2006; Koonin and Yutin, 2018), but it remains unclear where the root of the NCLDV should be placed, and other studies have recovered topologies that place the Asfarviruses as a sister group to other NCLDV families (Guglielmini et al.). For purposes of our analysis, here we kept GVMAG-S-3300005056-23 as a basal-branching Asfarvirus, but further studies are needed to confirm the evolutionary provenance of this MAG.

In addition to phylogenetic analysis, we also performed pairwise AAI analysis to assess the genomic divergence between different Asfarviruses. Our analysis recovered pairwise AAI values ranging from 27 to 75% (Figure 2) with mean and median values of 31.7% and 31.0% respectively. This result is consistent with the deep-branching clades identified in the phylogenetic analysis and confirms the high genomic divergence within the *Asfarviridae*.

**Figure 2.**
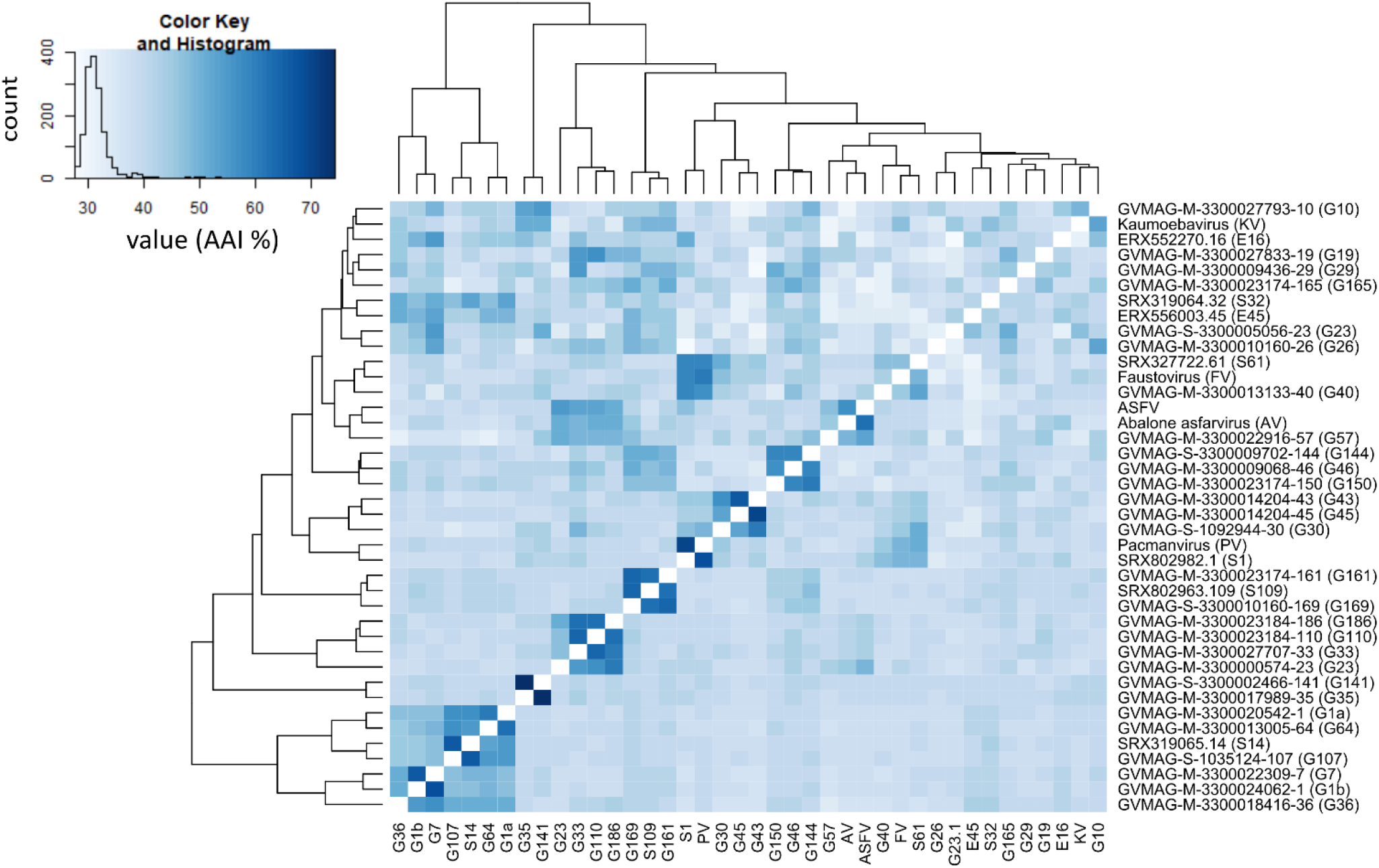
Amino acid identity percentage between the MAGs and reference Asfarviruses. The histogram inside the colorbar represents the frequency of AAI %.

### 3.3 Pan-genomics of the Asfarviruses

We found 7,410 total Orthologous groups (OGs), including 6,480 that were found in one Asfarvirus genome only. The number of unique OGs for each genome ranged from 48 to 428. We observed 12 core OGs in all viruses, including the major capsid protein (MCP), VLTF3-like transcription factor, A32 packaging ATPase, DNA topoisomerase II, DNA ligase, RNA polymerase subunit B, ATP dependent helicase hrpA, VVA8L-like transcription factor, and some hypothetical proteins (Figure 3). Because some of the Asfarvirus MAGs are incomplete, it is likely that the number of OGs conserved in all Asfarviruses is larger and includes other highly conserved proteins such as the B-family DNA Polymerase. Nonetheless, the high number of genome-specific OGs highlights the genomic diversity present in the *Asfarviridae* family, which is consistent with the high level of variability in other families of NCLDV (Van Etten et al., 2010b).

**Figure 3.**
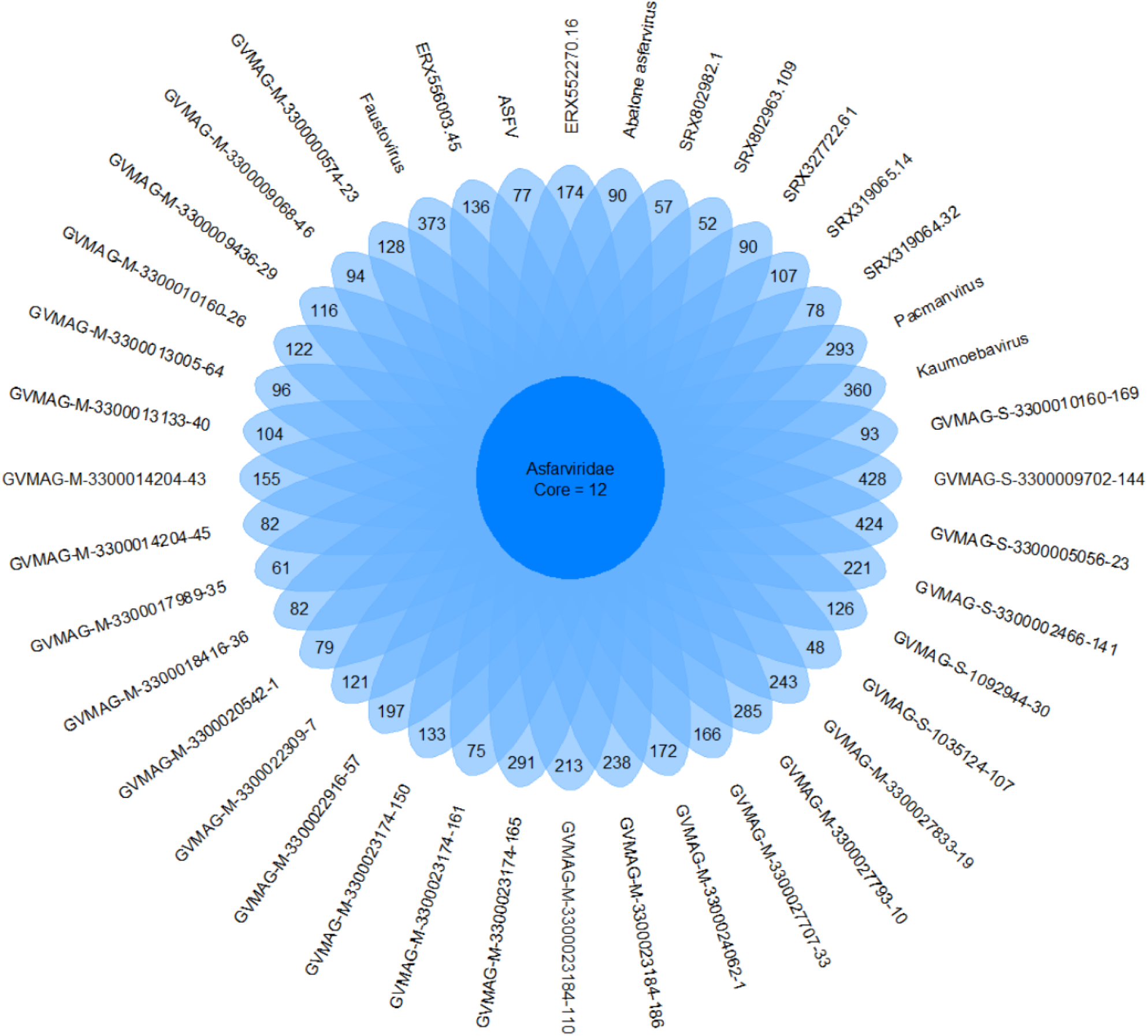
Unique and core genes shared between the MAGs and reference Asfarviruses. Here we define “core” as all genes found in 90% or more genomes.

In order to visualize the pattern of gene sharing, we performed bipartite network analysis using the Asfarvirus OGs, with six Poxvirus genomes used as non-Asfarvirus references. Given that virus evolution is characterized by extensive gene loss, gain, and exchange, this approach can be complementary to traditional phylogenetic analysis (Iranzo et al., 2016). The bipartite network showed some clustering of the MAGs based upon the habitat (Figure 4), although many co-clustered MAGs are also closely related and common gene content due to shared ancestry cannot be ruled out. The *Poxviridae* clustered separately in a small sub-network, indicating that their gene content is clearly distinct from the *Asfarviridae*. Hence, the bipartite network provides support for the phylogenetic findings we have for the Asfarviruses as well as depicts the gene- sharing pattern of these viruses.

**Figure 4.**
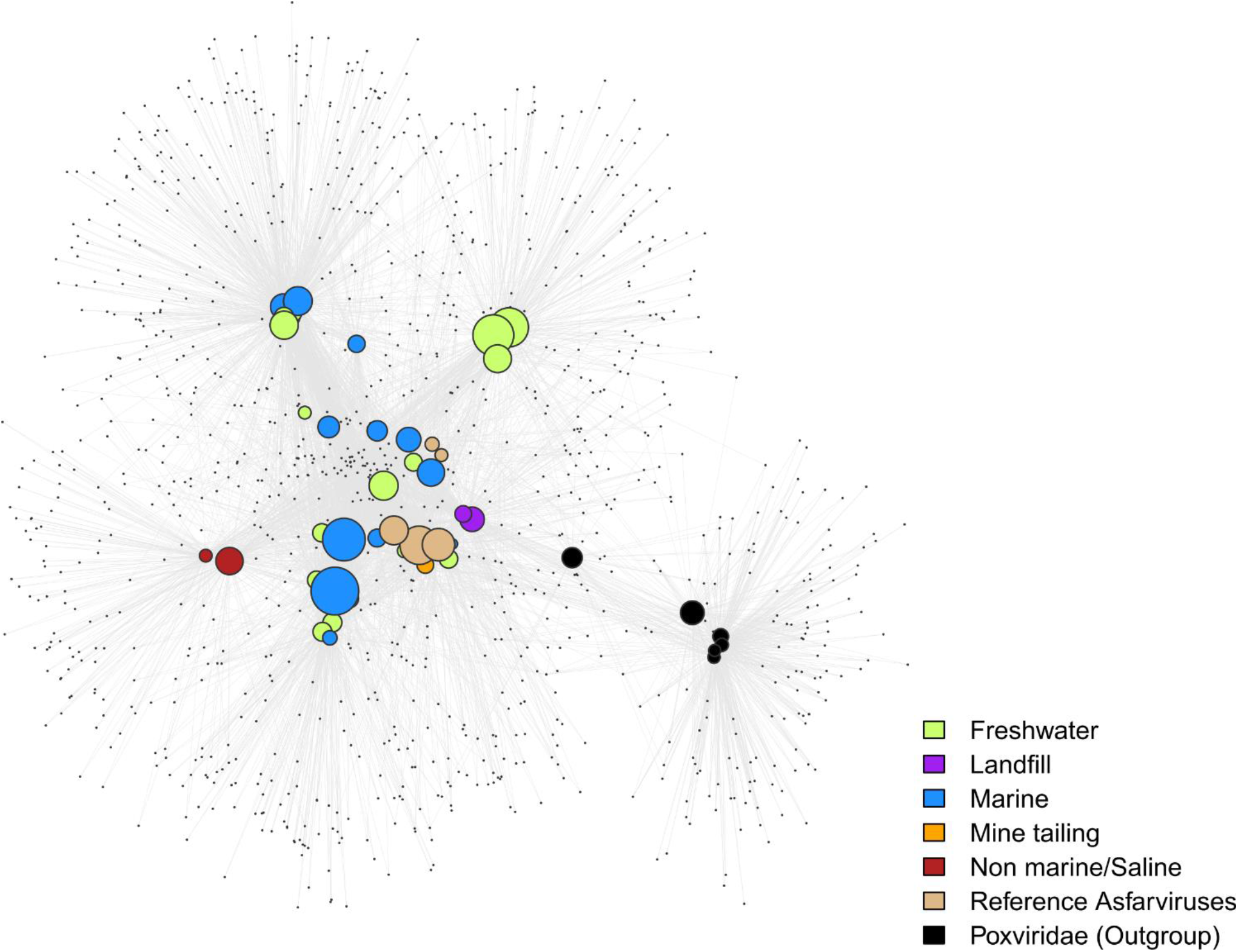
Bipartite network plot for the MAGs. The larger nodes represent genomes while the smaller nodes represent OGs/gene families. Genomes were connected to the genes if they contained one through edges and two or more genomes were connected through the gene family. MAGs are colored based on their habitat. The size of the larger nodes represents genome size.

### 3.4 Genomic Chimerism of the Asfarviruses

NCLDVs are known to have chimeric genomes with genes that are derived from multiple sources (Boyer et al., 2009), and we therefore sought to quantify the extent of this genomic chimerism in environmental Asfarviruses by comparing the encoded proteins of the Asfarvirus MAGs to the RefSeq database (see Methods for details; Supplementary Data 1). We found that between 40 to 70% of the proteins in each genome had no detectable hits to reference proteins, while 16-55% had best matches to other viruses, 5-22% to Eukaryotes, 3-15% to Bacteria, and 0-2% to Archaea (Figure 5A). We examined the proteins with best hits to Eukaryotes in more detail because this may provide some insight into host-virus gene exchange and therefore link these viruses to putative hosts. Overall, best hits to eukaryotes included matches to Animalia, Plantae, Fungi, and Protists such as Stramenopiles, Alveolata, Archaeplastida, Cryptista, Excavata, Choanomonada, Apusozoa, Porifera, and Amoebozoa (Figure 5B). The percent identity of these matches ranged from 19.4 - 93.2 (median 35.3), with only 4 greater than 90% identity suggesting that, if these represent gene exchanges between NCLDV and eukaryotes, the vast majority have not occurred recently. Although recent studies have revealed a dynamic gene exchange between NCLDV and eukaryotic lineages that can be used to link viruses to their hosts (Moniruzzaman et al., 2020b; Schulz et al., 2020), our analysis did not identify any clear signatures in the Asfarvirus MAGs that could be used for this purpose. It is possible that future work examining endogenous NCLDV signatures in eukaryotic genomes may be useful to better identify virus-host relationships.

**Figure 5.**
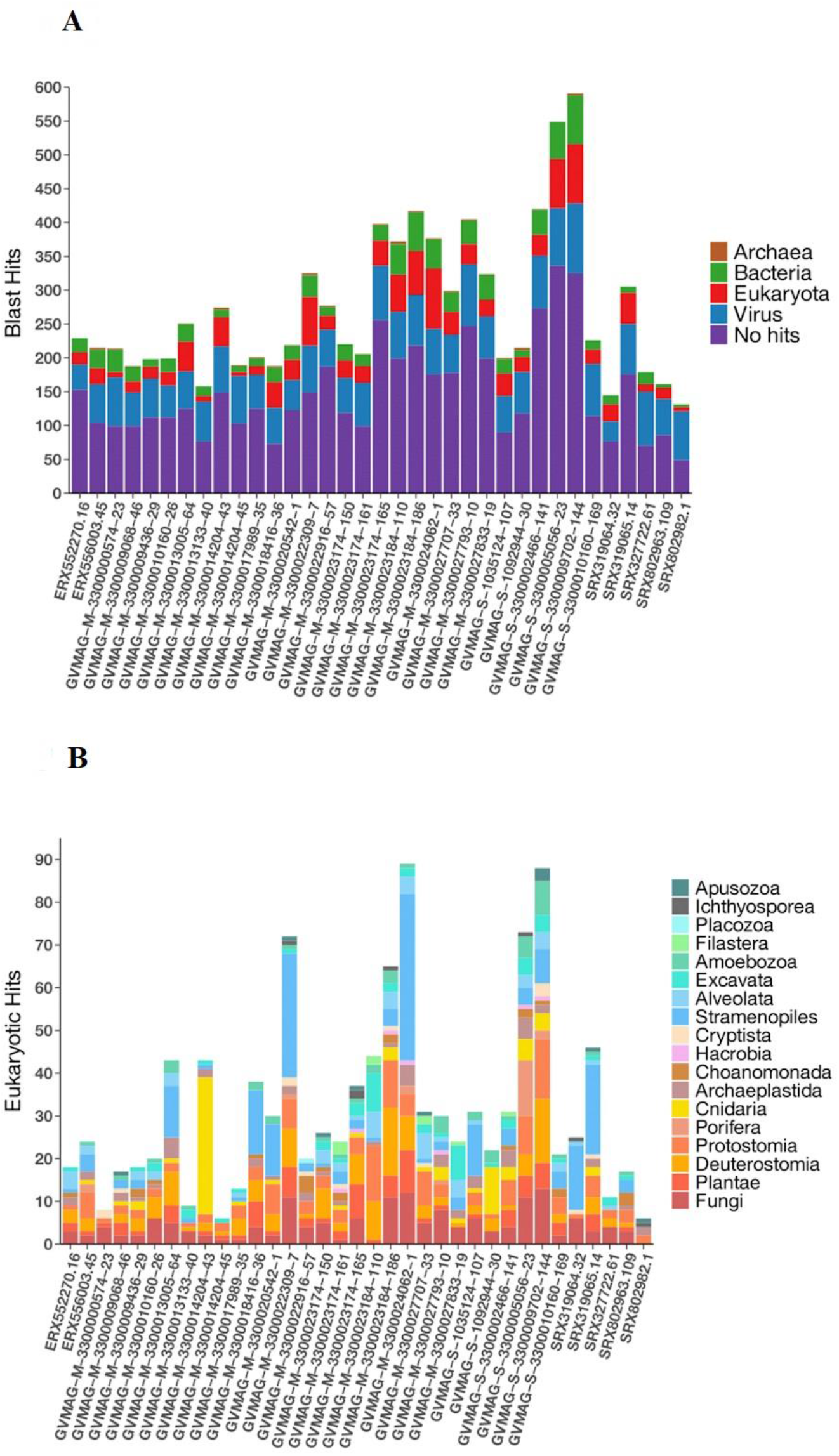
Distribution of homologous hits to MAGs determined by the BLASTp. **(A)** Total hits to three domains of life and viruses **(B)** eukaryotic hits.

### 3.5 Asfarvirus Genes Involved in Manipulating Host Metabolism

To assess the potential functions of the new MAGs proteins, we performed functional annotation using HMMER searches against the EggNOG database (all annotations available in Supplementary Data 2). As expected, in all MAGs we detected genes involved in DNA replication and repair, transcription, and post-translational modification, which is consistent with the prevalence of these functions across NCLDV (Yutin and Koonin, 2012) (Figure 6). Among the proteins involved in post-translational modification, we found genes responsible for ubiquitination (KOG0802 and KOG1812) and ubiquitin dependent proteins in 26 MAGs. Ubiquitination has been found to be an important counteracting mechanism to oxidative stress response in eukaryotes that direct the unwanted proteins to proteasome for degradation (Silva et al., 2015). In *Aureococcus anophagefferens* giant virus (AaV), ubiquitin dependent protein- ubiquitin ligases were found to be expressed within 5 minutes of virus-infection and is thought to be involved in degradation of host proteins (Moniruzzaman et al., 2018). The ubiquitin protein has also been reported in Marselliviruses, where it is thought to play an important role in host signaling (Boyer et al., 2009). A protein homologous to the ubiquitin-proteasome (UP) system has been found to be encoded by ASFV, suggesting its role during early infection and replication (Barrado-Gil et al., 2017). Hence, this suggests that ubiquitination may be a common mechanism across diverse Asfarviruses.

Genes predicted to be involved in carbohydrate metabolism were prevalent in the MAGs, consistent with previous findings that these genes are widespread in NCLDVs. We observed glycosyltransferase enzymes that are important in glycosylation of viral proteins in 15 Asfarvirus MAGs. These enzymes have been previously reported in giant viruses (Markine-Goriaynoff et al., 2004). Also, past studies have indicated the presence of glycosylating genes (Van Etten et al., 2010a; Piacente et al., 2015) and other enzymes involved in carbohydrate metabolism in NCLDVs (Fischer et al., 2010). Interestingly, we found genes involved in the shikimate pathway that is linked to the biosynthesis and metabolism of carbohydrates and aromatic amino acids (phenylalanine, tryptophan, and tyrosine) in five MAGs. We found 3-deoxy-7-phosphoheptulonate synthase (2QPSU) (the first enzyme in the shikimate pathway), chorismate synthase (KOG4492), and prephenate dehydrogenase (KOG2380) all in ERX556003.45 and only 3-deoxy-7-phosphoheptulonate synthase in four other MAGs. The shikimate pathway is widespread in bacteria, archaea, and protists but not in metazoans (Richards et al., 2006). We also found acetolactate synthase genes (KOG4166) in three MAGs. Acetolactate synthase that are involved in the synthesis of amino acids such as leucine, isoleucine, and valine has been previously described to be present in large DNA viruses infecting green algae mainly, *Prasinovirus* (Weynberg et al., 2009; Moreau et al., 2010; Zhang et al., 2015). Hence, the detection of these enzymes shows the potential role of the Asfarvirus MAGs in the manipulation of amino acid metabolism in their hosts during infection.

**Figure 6.**
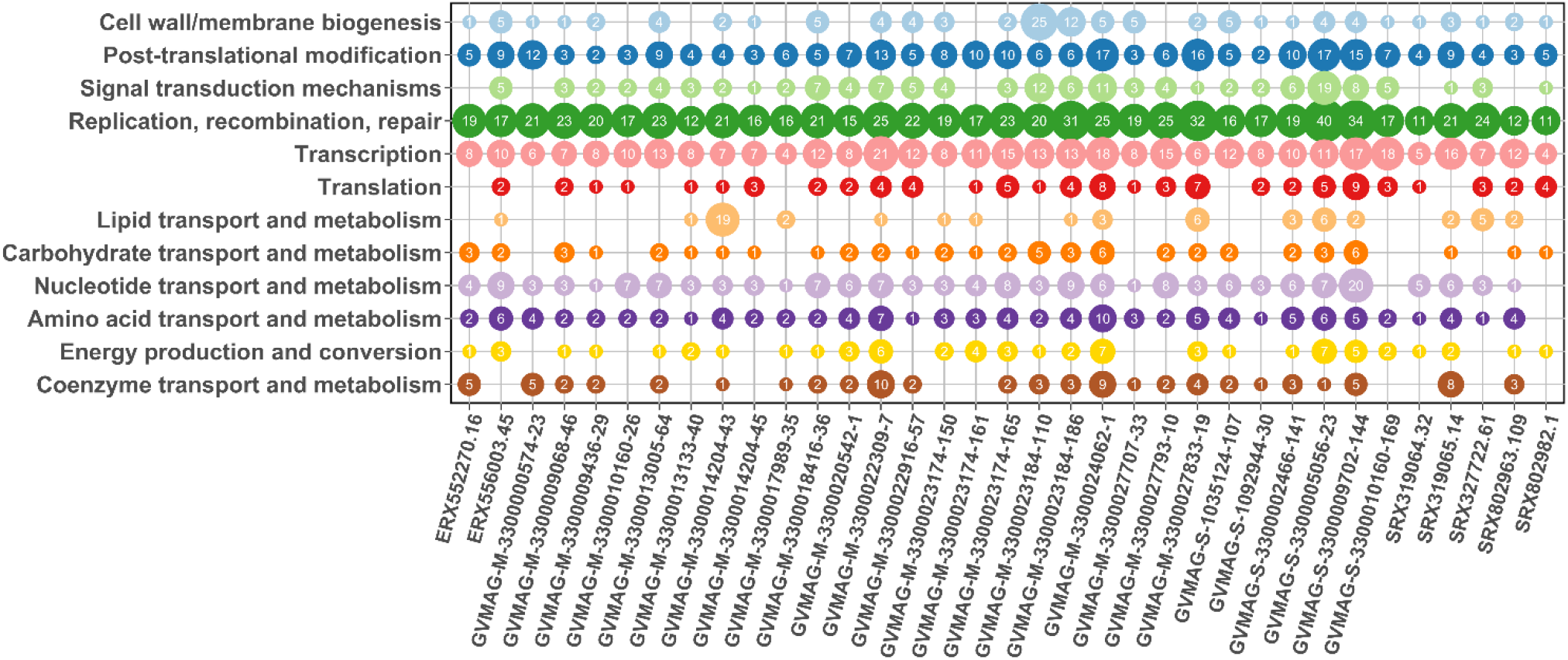
Protein annotation for MAGs. The x-axis represents the MAGs while y-axis represents the COG category. The number inside the bubble represents the number of genes present in that MAG that had the annotated function.

Genes responsible for signal transduction were also present in some of the MAGs. KOGs representing serine/threonine protein kinase and tyrosine/serine/threonine phosphatase were present in seven MAGS. These enzymes constitute a major form of signaling and regulation of many cellular pathways such as cell proliferation, differentiation, and cell death. Serine/threonine kinases have also been reported in Marseillevirus, Iridovirus, and Ascoviruses (Boyer et al., 2009; Piégu et al., 2015) and ASFV, suggesting that it might have a role in early infection and programmed cell death (apoptosis) (Baylis et al., 1993).

We found genes homologous to cysteine desulfurase (COG1104) proteins in 21 out of 35 MAGs (Supplementary Data 2). NifS genes whose presumed functions are similar to that of cysteine desulfurase are reported to be associated with ASFV, Faustovirus, and Pacmanvirus with possible involvement in host cell interactions (Andreani et al., 2017). Cysteine desulfurase proteins are found in bacteria and eukaryotes and are involved in the biosynthesis of iron (Fe) - sulphur (S) clusters, thiamine, biotin, lipoic acid, molybdopterin, NAD, and thionucleosides in tRNA (Mihara and Esaki, 2002). Hence, the discovery of the enzyme cysteine desulfurase adds to the viral proteins involved in electron transfer processes.

Gene encoding for cell redox homeostasis (KOG0191) and cellular response to nitrogen starvation (KOG1654) were also common among the MAGs. Nutrient limitation has the potential to reduce viral productivity; virus reproduction mostly depends upon the intracellular nitrogen and phosphorous pool during early infection while they might depend upon the extracellular nitrogen availability as infection proceeds (Zimmerman et al., 2020). Genes involved in responding to the nutrient starvation can influence the nutrient uptake and replication in these viruses. Overall, these results demonstrate that in addition to universal genes that play a role in host invasion and viral replication, Asfarviruses also contain genes involved in metabolism. Hence, capable of reprogramming the virocell metabolism (Moniruzzaman et al., 2020a).

### 3.6 Biogeography of marine Asfarviruses

While most cultured Asfarviruses were isolated from sewage samples, various metagenomic studies have revealed that NCLDVs are highly diverse and abundant in aquatic environments (Monier et al., 2008; Hingamp et al., 2013), and one recent study noted that Asfarviruses are prevalent in some marine samples (Endo et al., 2020). To examine the biogeography of the Asfarvirus MAGs in more detail we conducted a fragment recruitment analysis using reads from the Tara oceans expedition (Sunagawa et al., 2015). We examined 25 diverse metagenomic samples from surface and deep chlorophyll maxima (DCM) oceanic regions. The Asfarvirus MAG ERX552270.16 was present in five metagenomic samples, ERX556003.45 was found in 19, and GVMAG-M-3300027833-19 was found in one, revealing that some Asfarvirus are globally distributed in the ocean (Figure 7A). The fragment recruitment plots revealed that the MAGs had consistent coverage of reads with 100% nucleic acid identity matches to the metagenomic reads (Figure 7 B, C, D, and Supplementary Figures 1 & 2), demonstrating high similarity of these viruses across long distances. Few gaps were visible in the recruitment plots, indicating the absence of genomic islands in these viruses.

**Figure 7.**
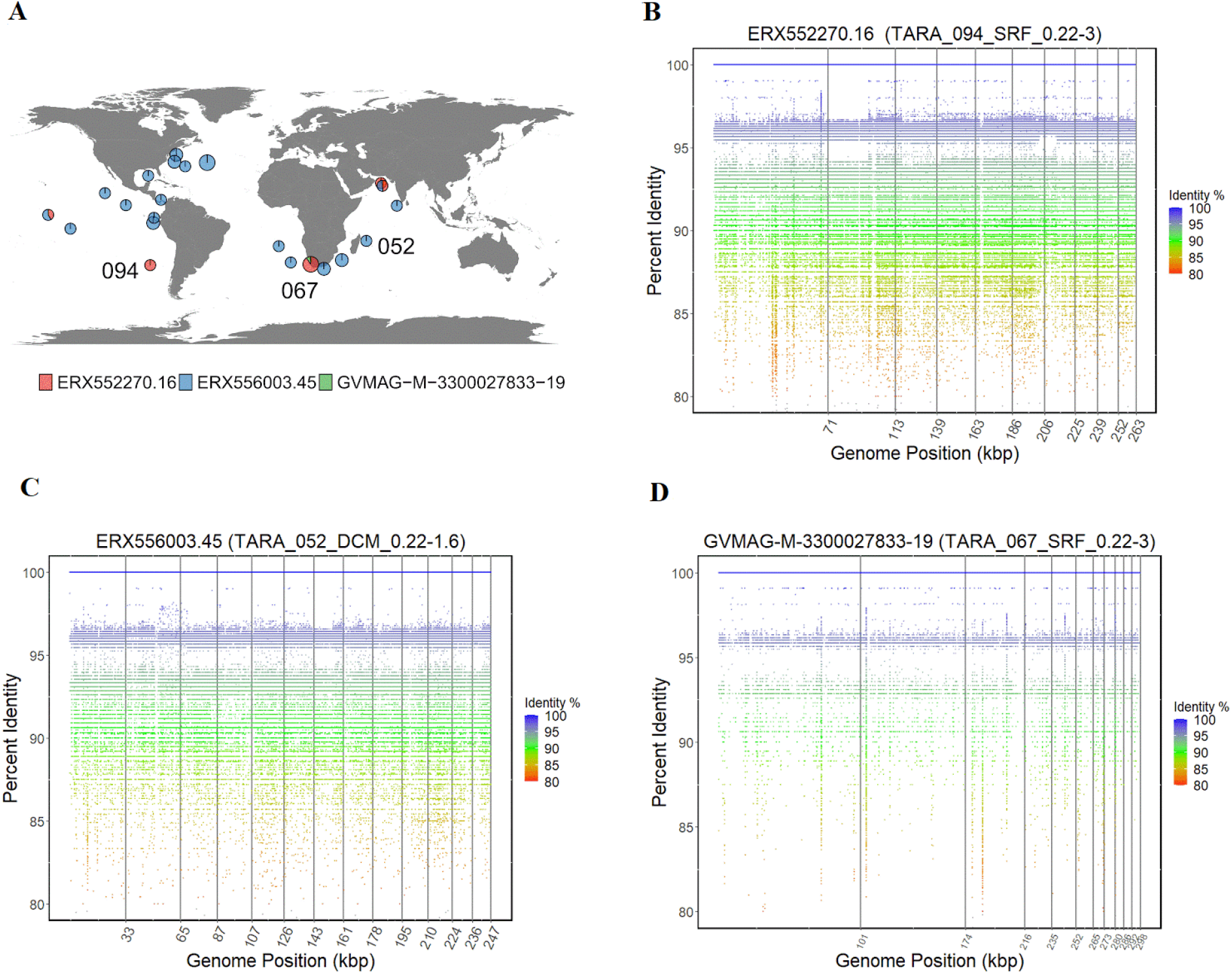
**(A)** Distribution of Asfarvirus matching metagenomic reads from the TARA ocean project. **(B)**, **(C)**, **(D)** Fragment recruitment plot for metagenomic reads to ERX552270.16, ERX556003.45, and GVMAG-M-3300027833-19 respectively. The x-axis of the recruitment plot shows position of the metagenomic reads along the genome length and y-axis represents the percent identity.

Previous studies have shown that the virus HcDNAV infects the marine dinoflagellate *Heterocapsa circularisquama,* which is responsible for harmful algal blooms in the marine environment. Although a complete genome of HcDNAV is not available, several marker genes from this virus have been sequenced, are available in NCBI and have been previously reported (Ogata et al., 2009).We found that the MAG ERX552270.16 bore high sequence similarity to the HcDNAV marker genes, indicating that this MAG represents a closely related virus that potentially infects the same host. The Family B Polymerase (YP_009507841.1), HNH endonuclease (YP_009507839.1), DNA directed RNA Polymerase (BAI48199.1), and DNA mismatch repair protein (mutS) (BAJ49801.1) of HcDNAV all had 95.8 to 99% amino acid identity to homologs in ERX552270.16 (Table 2). Moreover, we constructed a PolB phylogeny of the *Asfarviridae* that confirmed that these viruses cluster closely together (Figure 8). Lastly, our fragment recruitment analysis from Tara Ocean data confirmed that ERX552270.16 is widespread in the ocean (Figure 7 and Supplementary Figure 1), consistent with the hypothesis that it is a marine virus that also infects *Heterocapsa circularisquama* or a closely related dinoflagellate. Given these similarities, we anticipate that ERX552270.16 can be a useful reference genome for exploring the genomics and distribution of close relatives of HcDNAV.

**Table 2.** AAI between the HcDNAV genes (only genes available at NCBI) and MAG ERX552270.16 as analyzed by blastp.

| HcDNAV genes | MAG genes | AAI % |
| --- | --- | --- |
| HNH endonuclease (YP_009507839.1) | contig_19878_13 | 95.88 |
| DNA mismatch repair protein (BAJ49801.1) | contig_7191_19 | 98 |
| type B DNA polymerase (YP_009507841.1) | contig_19878_12 | 98.37 |
| DNA-directed RNA polymerase subunit (BAI48199.1) | contig_19878_28 | 98.97 |

**Figure 8.**
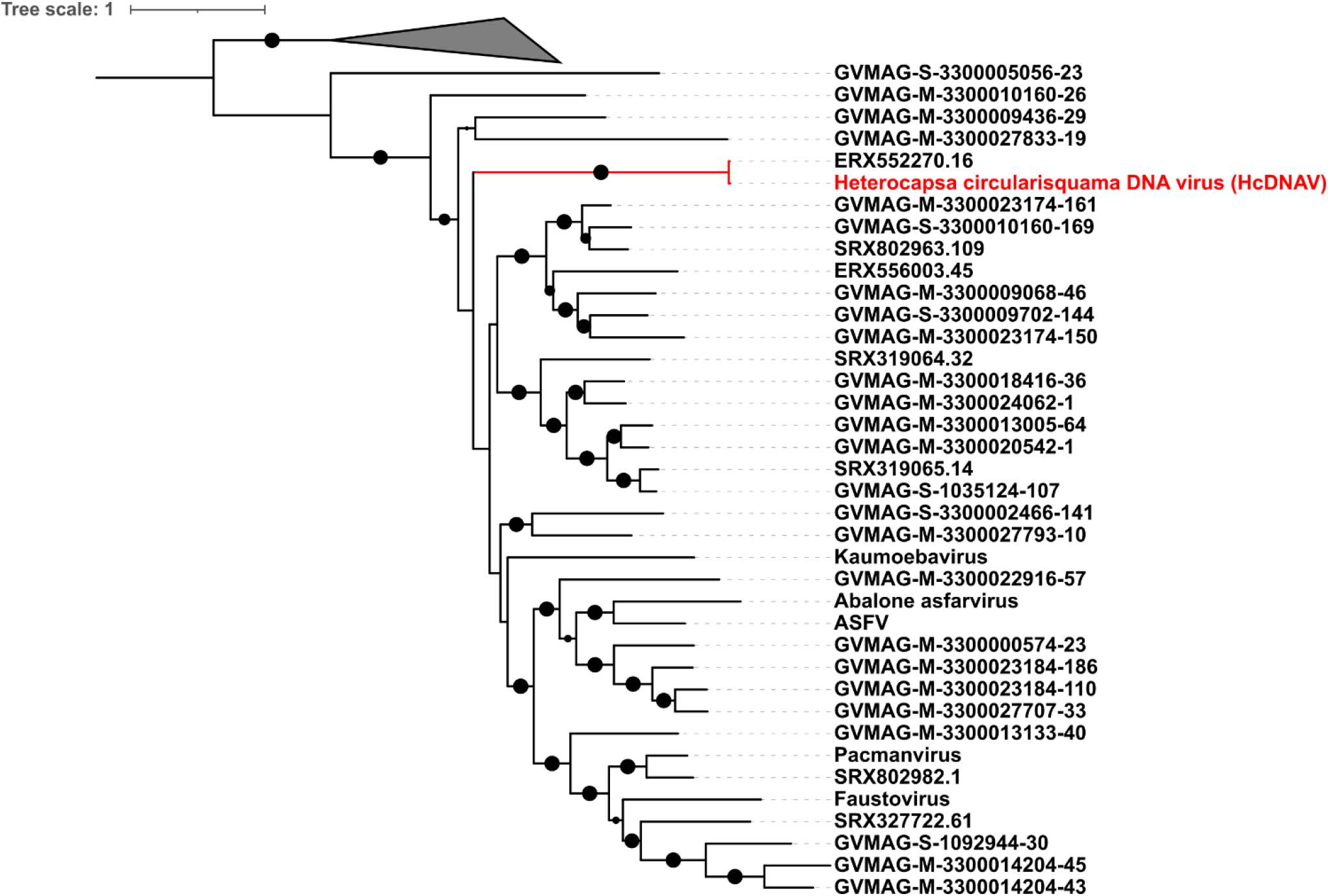
Phylogenetic tree reconstruction based on DNA polymerase B gene. (New reference virus HcDNAV has been added). The size of the black dot represents the bootstrap values. Only bootstrap values greater than 0.5 are shown.

## 4 Conclusion

While the ASFV was the only known member of *Asfarviridae* for many years, recent work has identified numerous additional members of this viral family. In this study, we leveraged the genomic characterization and phylogenetic relationships of *Asfarviridae* members and provide a robust phylogenetic and comparative genomic analysis of this viral family. Our results highlight the high level of genomic and phylogenetic divergence between disparate members of the *Asfarviridae*, and homology searches suggest that many genes within this viral group are the product of ancient horizontal transfers from cellular lineages. Moreover, we provide fragment recruitment plots that confirm that some Asfarviruses are ubiquitous in the ocean, where they may infect ecologically important protists such as bloom forming dinoflagellates. These findings suggest that diverse Asfarviruses are broadly distributed in the environment and play important roles in numerous ecosystems.

## Conflict of Interest

The authors declare no competing interest.

## Author Contributions

FOA designed the study, SK and MM performed the experiment, and SK and FOA wrote the manuscript.

## Funding

This research was funded by a Simons Foundation Early Career Award in Marine Microbial Ecology and Evolution and an NSF IIBR award 1918271 to FOA.

## Acknowledgments

We acknowledge the use of the Virginia Tech Advanced Research Computing Center for bioinformatic analyses performed in this study. We are thankful to the members of Aylward lab for their helpful suggestions.

## Data Availability Statement

The datasets analyzed in this study are already publicly available and were accessed as described in the Methods section.

## Supplementary Materials

All Supplementary informations are provided in the Supplemtary materials file.

## Notes

### Competing Interest Statement

The authors have declared no competing interest.

