## Supplementary material for "Comparative Genomics and Environmental Distribution of Large dsDNA viruses in the family *Asfarviridae*": Supplementary material.pdf

### 1 Supplementary Data

**Supplementary Data 1**. Best hits to the MAGs as determined by blastp.

**Supplementary Data 2**. Protein annotation of the viral MAGs.

### 2 Supplementary Figures

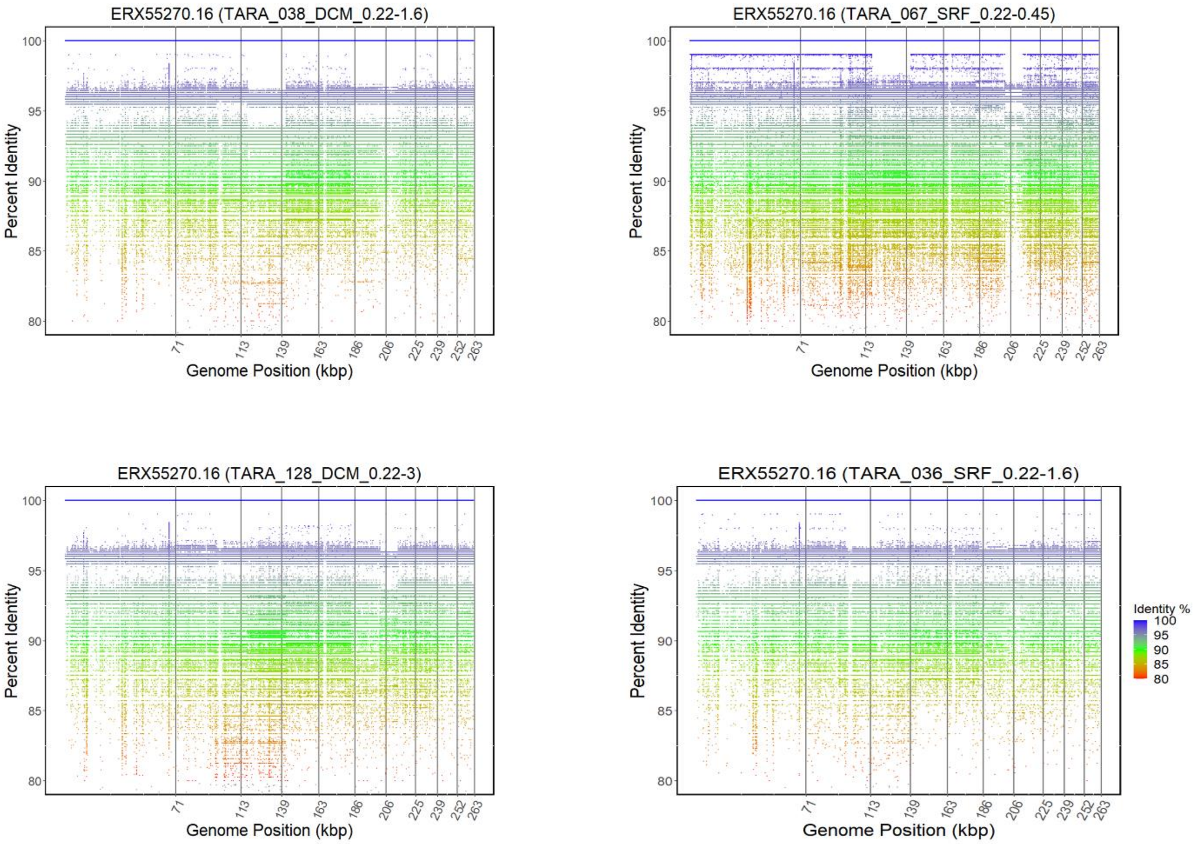


**Supplementary Figure 1.** Fragment recruitment plot for metagenomic reads to ERX552270.16. The x-axis of the recruitment plot shows position of the metagenomic reads along the genome length and y-axis represents the percent identity.

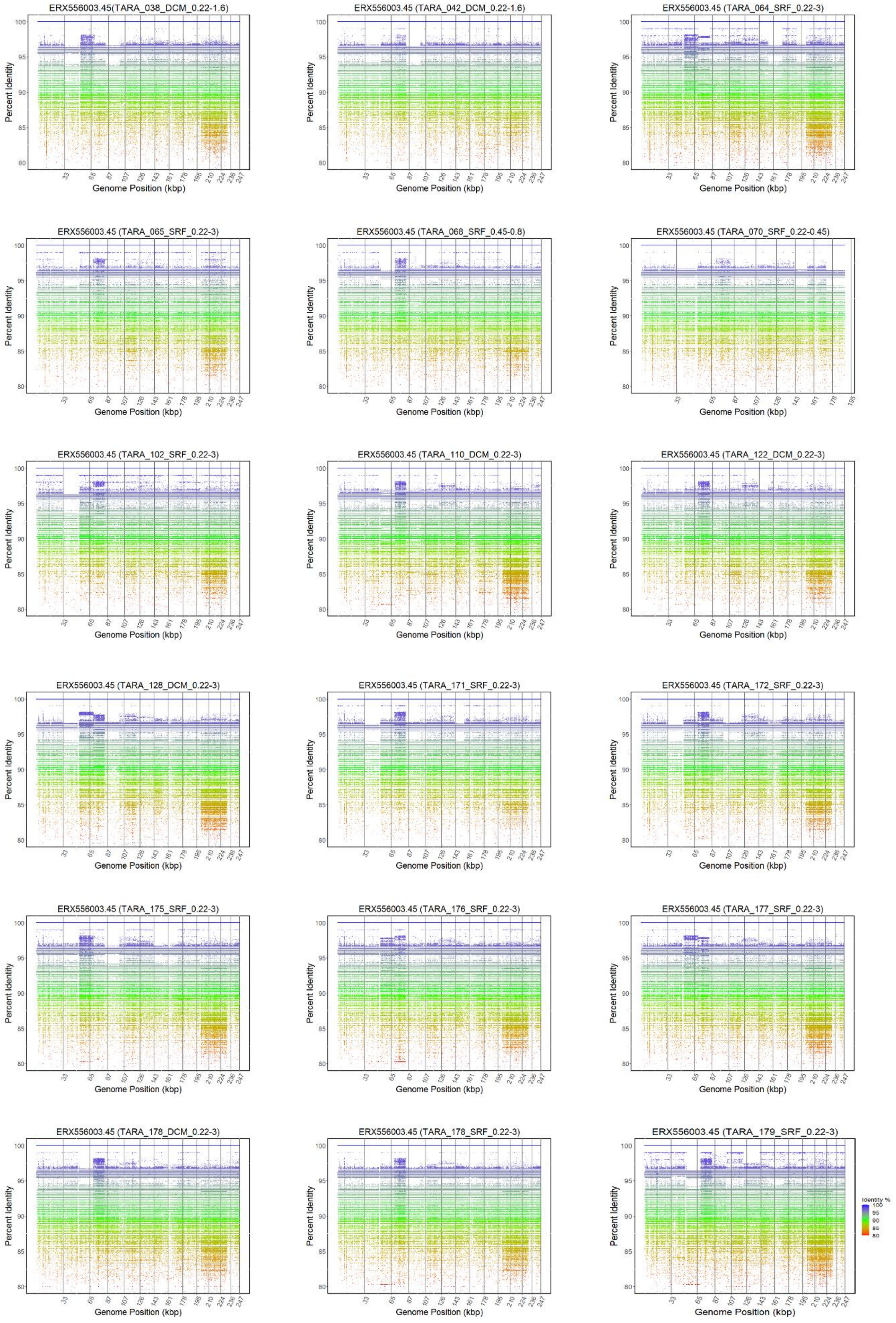
ERX556003.45(TARA_038_DCM_0.22-1.6)
ERX556003.45 (TARA_042_DCM_0.22-1.6)
ERX556003.45 (TARA_064_SRF_0.22-3)
ERX556003.45 (TARA_065_SRF_0.22-3)
ERX556003.45 (TARA_068_SRF_0.45-0.8)
ERX556003.45 (TARA_070_SRF_0.22-0.45)
ERX556003.45 (TARA_102_SRF_0.22-3)
ERX556003.45 (TARA_110_DCM_0.22-3)
ERX556003.45 (TARA_122_DCM_0.22-3)
ERX556003.45 (TARA_128_DCM_0.22-3)
ERX556003.45 (TARA_171_SRF_0.22-3)
ERX556003.45 (TARA_172_SRF_0.22-3)
ERX556003.45 (TARA_175_SRF_0.22-3)
ERX556003.45 (TARA_176_SRF_0.22-3)
ERX556003.45 (TARA_177_SRF_0.22-3)
ERX556003.45 (TARA_178_DCM_0.22-3)
ERX556003.45 (TARA_178_SRF_0.22-3)
ERX556003.45 (TARA_179_SRF_0.22-3)
Percent Identity
Genome Position (kbp)
Identity %
100
95
90
85
80

**Supplementary Figure 2.** Fragment recruitment plot for metagenomic reads to ERX556003.45. The x-axis of the recruitment plot shows position of the metagenomic reads along the genome length and y-axis represents the percent identity.
